# *CircPLXNA2* affects the proliferation and apoptosis of myoblast through *circPLXNA2/gga-miR-12207-5P/MDM4* axis

**DOI:** 10.1101/2023.01.01.522409

**Authors:** Xu Dong, Jia-bao Xing, Qingchun Liu, Mao Ye, Zhen Zhou, Yantao Li, Zhenhui Li, Qinghua Nie

## Abstract

**Background:** circRNAs are new-identified special endogenous RNA molecules that covalently close a loop by backsplicing with pre-mRNA.

In the cytoplasm, circRNAs would act as molecular sponges to bind with specific miRNA to promote the expression of target genes. However, there is still in its fancy of knowing circRNA functional alternation in skeletal myogenesis. In this study, we favor a model to identify the circRNA-miRNA-mRNA interaction network in which the axis may be implicated in the progression of chicken primary myoblasts (CPMs) myogenesis by a combination of multi-omics (i.e., circRNA-seq and ribo-seq).

**Results:** In total, 314 circRNA-miRNA-mRNA regulatory axis containing 66 circRNAs, 70 miRNAs, and 24 mRNAs that may be relevant to myogenesis were collected. With these, the *circPLXNA2-gga-miR-12207-5P-MDM4* axis aroused our research interest. The *circPLXNA2* is highly differentially expressed during differentiation versus proliferation. It was demonstrated that *circPLXNA2* inhibited the process of apoptosis while at the same time stimulating cell proliferation.

Furthermore, we demonstrated that *circPLXNA2* could inhibit the repression of *gga-miR-12207-5p* to *MDM4* by directing binding to *gga-miR-12207-5p*, thereby restoring *MDM4* expression.

**Conclusions:** *CircPLXNA2* could function as a competing endogenous RNA (ceRNA) to inhibit the repression of *gga-miR-12207-5p* to *MDM4* by directing binding to *gga-miR-12207-5p*, thereby recovering the expression of *MDM4*.

## Background

Chicken is an economically domestic animal that plays an indispensable role in human meat consumption as the first-consumed food source. Despite its economic value, the muscle growth and development of chicken are of great importance, and have become the industry’s primary focus. Skeletal muscle, as an essential component of the body, plays a crucial role in maintaining movement, regulating metabolism, etc. Importantly, the growth and development of skeletal muscle are closely related to meat yield [1]. Multiple studies have demonstrated in recent years that the non-coding RNA (ncRNA) family regulates the growth and development of skeletal muscle to a large extent, due to the genome’s predominance (more than 98%) [2].

Among these non-coding RNA molecules, circRNA is a newly discovered type of endogenous RNA that forms a closed-loop structure when pre-mRNA is covalently spliced into it via back-splicing [2, 3]. It is common knowledge that circRNA possesses a distinctive biological structure, and its location determines the multiple regulatory functions it can perform [4]. CircRNA is resistant to RNase R digestion and has a high stability because it lacks a 5’cap and 3’ploy A tail [5]. CircRNA sequencing platform had been successfully deployed, and a significant amount of circRNA had been characterized since its inception [6, 7]. Using this sequencing method, several circRNAs have been identified as associated with the muscle growth process, such as *CircZNF609* and *CircLMO7* [8, 9]. However, it is for such a short time span of discovery that there is still in its fancy of knowing circRNA functional alternation in skeletal myogenesis.

MicroRNA (miRNA) is another class of endogenous non-coding RNA molecules that regulates biological phenotype by target binding 3’UTR of mRNA. Since the discovery of the very first microRNA in 1993, research into the biological role of miRNA has covered a wide range of topics [10]. Numerous studies have shown that miRNA is very important in controlling the skeletal muscle phenotype. *MiR-27b-3p*, for instance, is suggested to exert a particular effect on muscle atrophy through *Cbl-b* gene [11]. For another, *miR-1*, *miR-133*, and *miR-499* have also been reported to regulate myogenesis through binding to 3’UTR of mRNA [12, 13]. However, a particular molecular sponge known as competing endogenous RNA may be able to reverse the binding effect (ceRNA, i.e., circRNA).

*P53* gene is the most inactivated tumor inhibitor gene in cancer [14–16]. The function of *p53* may be activated as a result of the heterogeneous pressure in one of two ways: cell cycle arrest or apoptosis [17, 18]. Several factors rarely activate the *p53* gene to ensure normal cell cycle under physiological conditions [19]. Murine Double Minute (MDM) 4, an important *p53*-binding protein, could interact with *MDM2* to form *MDM2/MDMX* heterodimer to act as an important negative regulator of *p53* upstream signaling pathway [20]. Numerous studies have demonstrated its biological role to regulate cell growth, but only a few have identified its potential role of co-regulating myogenesis with circRNA [14, 15, 20, 21].

In this study, we combined circRNA sequencing and ribosome footprinting to characterize a potential interaction network involving circRNA, miRNA, and mRNA that may regulate muscle development. In this profiling, we discovered a new circRNA called *circPLXNA2*, which we found to have the potential to interact with the molecules *gga-miR-12207-5p* and *MDM4*. We used functional assays to investigate the mechanism of phenotype alternation and potential interactions.

## Materials and methods

### Animals and Cell

Thigh muscles of 11-embryo aged chickens were isolated on a sterile forceps using autoclaved surgical equipment and then completely minced. The crushed muscle tissue was digested in a constant temperature incubator at 37°C for 15min using 0.25% trypsin-EDTA (Gibco, USA). Cells were treated with high glucose Dulbecco’s Modified Eagle medium (Gibico) containing fetal bovine serum (10% (v/v) fetal bovine serum (FBS) (pan) and 0.2% penicillin/streptomycin) to terminate digestion. The digested suspension was then filtered through a 70 μm strainer and centrifuged at 1000g for 5min. The precipitated cells were seeded uniformly in cell culture dishes by blowing with complete medium, and adherent purification was performed twice for 40min at 37□ in an incubator with 5% CO_2_ to eliminate the interference of chick embryo fibroblasts. Finally, the medium containing CPM cells was uniformly seeded in cell culture dishes for subsequent experiments.

### Parallel generation of CircRNA and riboRNA libraries

We prepared the CPM samples at growth medium (GM) and differentiation medium (DM) each for three biological replicates with good cell viability, respectively. Each sample were individually pooled for subsequent RNA extraction and library construction. Total RNA was extracted using TRI-zol®Reagent (Invitrogen) according to manufacturers’ instruction. Subsequently, the total RNA was treated with RNase R to degrade the linear RNAs, and purified using RNeasy MinElute Cleanup Kit (Qiagen, Germany). Genomic DNA was then removed using DNase I (Takara, Japan). CircRNA libraries were constructed using the TruSeqTM Stranded Total RNA Sample Preparation Kit from Illumina (Illumina, USA) after removal of host ribosomal RNA using Illumina Ribo-ZeroTM rRNA Removal Kits (Qiagen) and RNase R treatment. Next, strand-specific library was constructed using VAHTS Total RNA-seq (H/M/R) Library Prep Kit (Vazyme, Nanjing, China) for Illumina following the manufacturer’s instructions. Briefly, ribosome RNAs were removed to retain circRNAs. The enriched circRNAs were fragmented into short fragments by using fragmentation buffer and reverse transcribed into cDNA with random primers. Second-strand cDNA were synthesized by DNA polymerase I, RNase H, dNTP (dUTP instead of dTTP) and buffer. Next, the cDNA fragments were purified with VAHTSTM DNA Clean Beads (Vazyme), end repaired, poly(A) added, and ligated to Illumina sequencing adapters. Then UNG (Uracil-N-Glycosylase) was used to digest the second-strand cDNA. The digested products were purified with VAHTSTM DNA Clean Beads, PCR amplified, and sequenced using Illumina Novaseq 6000 (Illumina).

### Ribosome footprinting

Analogously, we prepared the CPMs during GM and DM each for three as biological replicates. The ribosome footprints were generated after immunoprecipitation of cardiomyocyte-specific monosomes with anti-HA magnetic beads after treating the lysate with RNase I. Then the libraries were constructed according to the manufacturer’s instruction using mammalian ribo-seq kit (Illumina) with each replicate. And the libraries were sequenced using Illumina Novaseq 6000 (Illumina).

### CircRNA Validation

We first validated the biological structure of *circPLXNA2* using convergent and divergent primers. Then the junction sequence of the *circPLXNA2* was sequenced by Tsingke Biotechnology Co., Ltd. (Beijing, China). Besides, we detected the RNase R sensitivity using RT-qPCR.

### Primers

All primers for this study were designed in Premier Primer 5.0 software (Premier Biosoft International), and synthesized by TSINGKE Biology Co (Guangzhou, China). The detailed information of all primers is listed in Table 1.

**Table 1.**
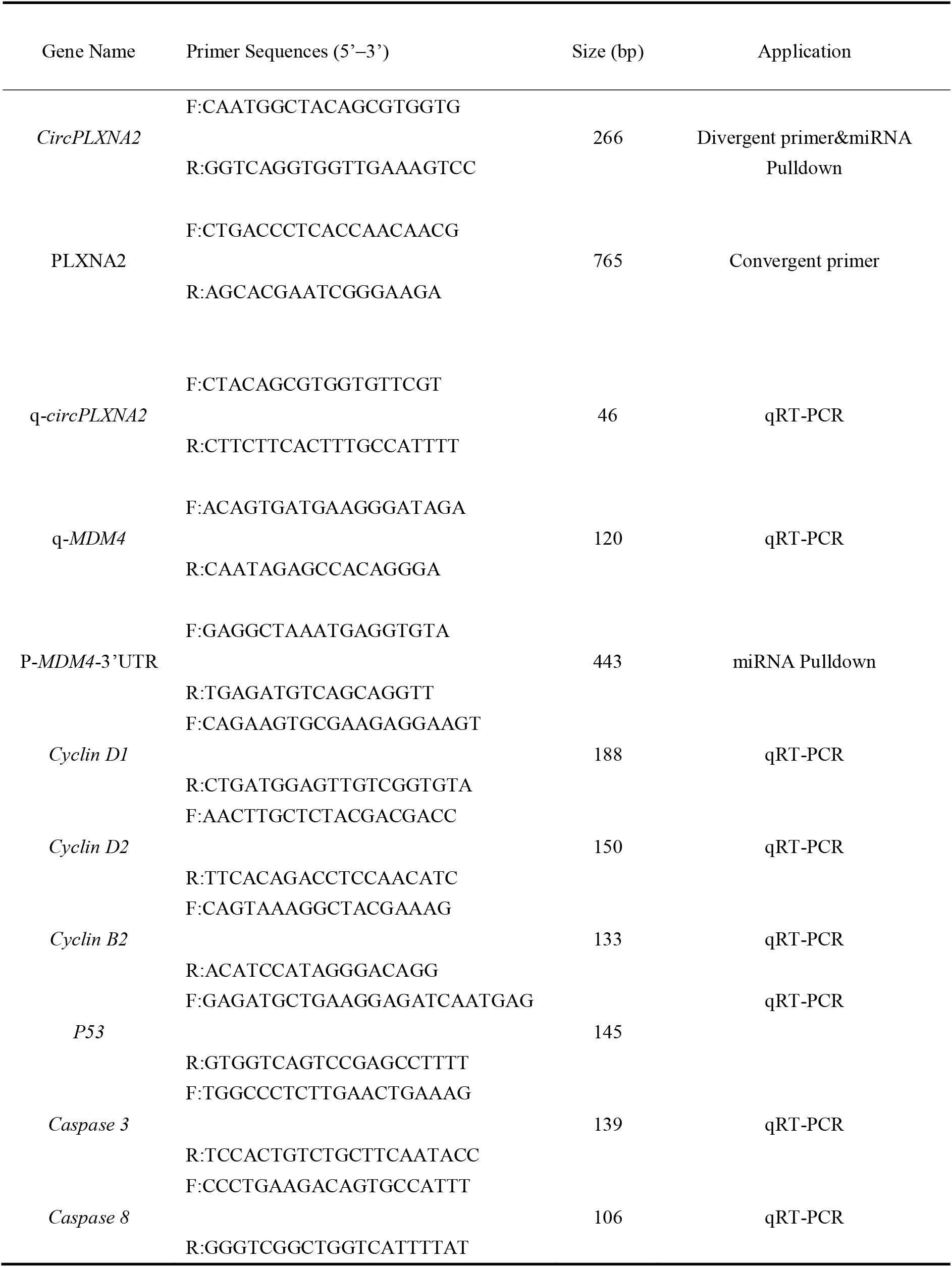
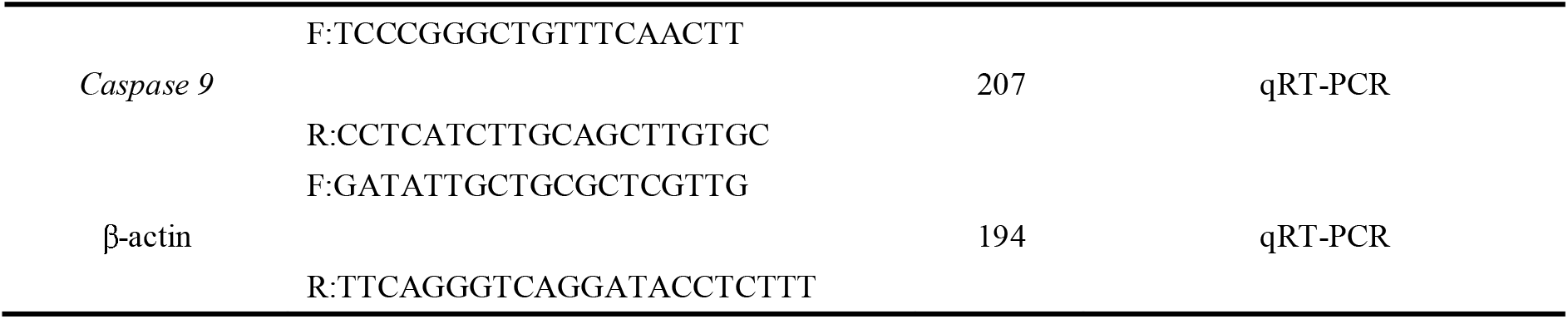
Primers used in this study.

**Table 2.**
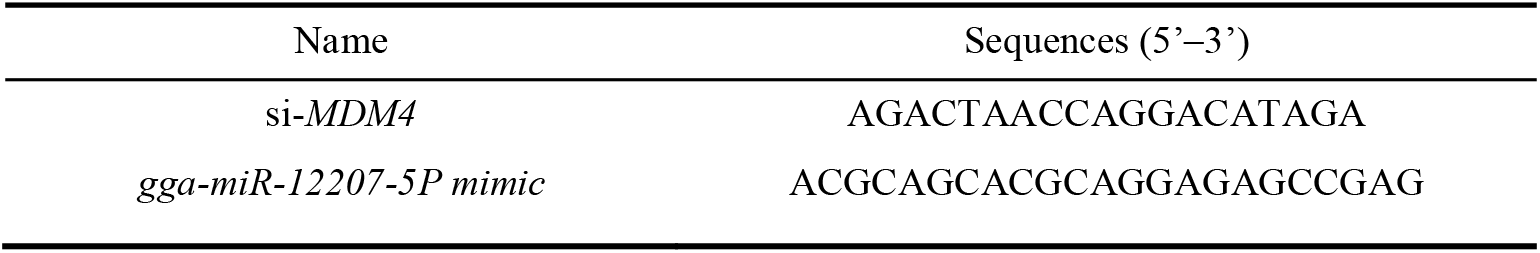
Oligonucleotide sequences in this study.

### RNA oligonucleotides and plasmids construction

*Gga-miR-12207-5P mimic*, *mimic NC*; *si-MDM4* and siRNA negative control were designed and synthesized by RiboBio (Guangzhou, China).

Construction of the *MDM4* overexpression vector. The full-length sequence of *MDM4* was amplified by conventional PCR primers, and the amplified sequence fragment was cloned into the commercial vector pcDNA3.1 (Promega, Madison, WI, USA) through the restriction site of HindIII and XhoII.

Construction of the *circPLXNA2* overexpression vector. First, the full-length linear sequence of *circPLXNA2* was synthesized according to the sequencing results. Subsequently, EcoRI restriction site, forward looping mediate sequence, and AG receptor were added into the 5’end of the forward primer. Similarly, the BamHI restriction site, reverse cyclization mediating sequence, and GT donor were added into the 5’end of the reverse primer. A homology arm was then added to the full-length linear sequence of *circPLXNA2* by PCR reaction, and the fragment was cloned into the circular vector pCD25-ciR by homologous recombinase (Geneseed Biotech, Guangzhou, China).

Construction of PmirGLO dual-luciferase target reporter vector. The segment sequences of *circPLXNA2* and *MDM4* 3’UTR containing the putative *gga-miR-12207-5P* binding sequence were amplified by PCR reaction and then subcloned into the XhoI and SalI restriction sites in the pmirGLO dual-luciferase reporter vector (Promega, USA).

### Cell transfection

All cell transient transfections in this study were performed at cells grew to 60-80% according to the manufacturer’s protocol of Lipofec-tamine 3000 reagent (Invitrogen). The nucleic acids were also diluted using OPTI-MEM medium (Gibco). For miRNA mimic, *mimic NC*, and siRNA transfection, the final transfection concentration was 100nM.

### Dual-luciferase reporter assay

In order to verify the binding relationship between *gga-miR-12207-5P* and *circPLXNA2* and *MDM4*, we designed the constructed plasmid for several groups of co-transfection experiments, i.e., a. *gga-miR-12207-5P mimic+Pmir-GLO-circPLXNA2-WT*, b. *mimic NC+Pmir-GLO-circPLXNA2-WT*, c. *gga-miR-12207-5P mimic+Pmir-GLO-circPLXNA2-MUT*, d. *mimic NC+Pmir-GLO-circPLXNA2-MUT*, e. *gga-miR-12207-5P mimic+Pmir-GLO-MDM4-WT*, f. *mimic NC+Pmir-GLO-MDM4-WT*, g. *gga-miR-12207-5P mimic+Pmir-GLO-MDM4-MUT*, and h. *mimic NC+Pmir-GLO-MDM4-MUT* were transfected into DF-1 cell lines with 60-80% confluence (96-well plate). After 48 h post transfection, luciferase activity analysis was examined using Fluorescence/Multi-Detection Microplate Reader (BioTek, Winooski, USA) and Dual-GLO^®^Luciferase Assay System Kit (Promega). Firefly luciferase activities were normalized to Renilla luminescence in each well.

### 5-Ethynyl-20-Deoxyuridine (EdU) Assays

To investigate the proliferation of the cells, 50μM 5-ethynyl-20-deoxyuridine was first added into cell to incubated for 2h after 36h transfection (EdU) (RiboBio, China), with fixing with 4% paraformaldehyde (PFA) for 30 min and neutralizing with 4%PFA with 2 mg/mL glycine, and then permeabilized with 0.5%Triton X-100. Cells were subsequently stained using the C10310 EdU Apollo In Vitro Imaging Kit (RiboBio). Finally, fluorescence microscopy (DMi8; Leica, Germany) was used to obtain cell images. At least 3 random fields were selected for each well to photograph, and Image J was used to count the cells, hence the cell proliferation rate was calculated.

### Biotin-Coupled miRNA Pull Down Assay

When myoblasts in two 10-cm dishes reached 60-80% confluence, 3’end biotin-labeled *gga-miR-12207-5P mimic* or *mimic NC* (100 nM, P < 0.05) was transfected into the cells alternatively. Cells were collected and washed with PBS after 36h transfection. Using miRNA pulldown Kit (BersinBio, China), the lysed cells were incubated with the sealed magnetic beads for 4 hours at 4 ° C with mild rotation (20 rpm/min) to pulldown the biotin-coupled RNA complex. The RNA was eluted and precipitated as required to obtain the RNA for subsequent RT-qPCR that specifically interacted with *gga-miR-12207-5P*.

### RNA Extraction, cDNA Synthesis and Quantitative Real-Time PCR

As described, total RNA was extracted from tissues or cells using TRIzol reagent^®^(Invitrogen) according to the manufacturer’s requirements. The concentration and integrity of extracted RNA were tested using NanoDrop One and 1% gel electrophoresis. Then the cDNA was synthesized using PrimeScript RT Reagent Kit with gDNA Eraser (Perfect Real Time) (TaKaRa). Real-time quantitative PCR was performed as previously described [13, 22, 23]

### Western blotting assay

The cells were incubated with RIPA buffer containing protease inhibitors (protease inhibitors: RIPA=1:100, (Solarbio, China) for 15min on ice to fully lyse the cells 48h after transfection. Then the cells were centrifuged at 12000 g for 10min at 4 °C, and the supernatant was removed. Total protein was then quantified using the BCA Protein Assay Kit (Beyotime, Shanghai, China). Proteins were separated in 12%SDS-PAGE and transferred to nitrocellulose membranes (Whatman, UK), blocked with 5% skim milk powder for 1 h, and then incubated with primary antibody solution overnight at 4°C. Subsequently, PVDF membranes were washed three times for 5 min with TBST solution (Beyotime) and then incubated with secondary antibody solution for 60 min at room temperature. Western immunoblotting results were analyzed using the Odyssey Fc system (LI-COR, Lincoln, NE, USA). Antibody information is as follows: Actived-Caspase-3 p17 polyclonal antibody (BS7004; Bioworld, USA; 1: 500) Cleaved Caspase-8 (Asp391) (18C8) Rabbit mAb (9496; Cell Sig-naling Technology, USA; 1:1000), Anti-Caspase-9 anti-body [E23] (ab32539; Abcam, UK; 1:1000), rabbit anti-GAPDH (AB-P-R 001; Hangzhou Goodhere Biotechnology Co. Ltd., China; 1:1,500).

### Statistical analysis

In this study, all results were showed as mean±S.E.M with 3-6 independent replications. An independent sample t-test was used to test the statistically significant difference between groups. We considered p < 0.05 to be statistically significant. * p < 0.05; ** p < 0.01; *** p < 0.001.

## Results

### The circRNA-miRNA-mRNA axes network is constructed by multi-omics

We used circRNA-seq and ribo-seq on chicken primary myoblasts (CPMs) to identify candidate circRNA involved in muscle growth regulation by examining differentially expressed circRNA and regulatory genes. First, circRNA-seq identified 14241 circRNAs with a centralized length span of 0-1000 bp, of which 360 were significantly differentially expressed. (Fig1 A-C). After filteration, 200 circRNAs were identified as up-regulating, and 160 circRNAs were identified as down-regulating in DM relative to GM of CPMs (Fig1 B-C). We mapped differentially expressed circRNAs to their parental genes and performed GO and KEGG enrichment analysis. GO enrichment analysis suggested a large variety of functional genes enrichment in the process of cellular development and differentiation, and KEGG biological pathway identification indicated the genes were involved in the signal pathway such as *MAPK*, *Wnt*, and *autophagy* (Fig 1 D-E). In addition, a total of 3342 differentially expressed genes (DEGs) were identified by ribo-seq profile, in which 2185 DEGs were up-regulating expressed, and 1157 DEGs were down-regulating expressed (Fig 1 F-G). GO enrichment analysis enriched the most genes into the biological process such as biological regulation, signaling, cell proliferation. KEGG signaling pathway analysis indicated that most signaling pathways associating with muscle development were regulated such as *p53*, *MAPK*, *mTOR*, and *JAK-STAT* (Fig 1 H-I).

**Fig.1.**
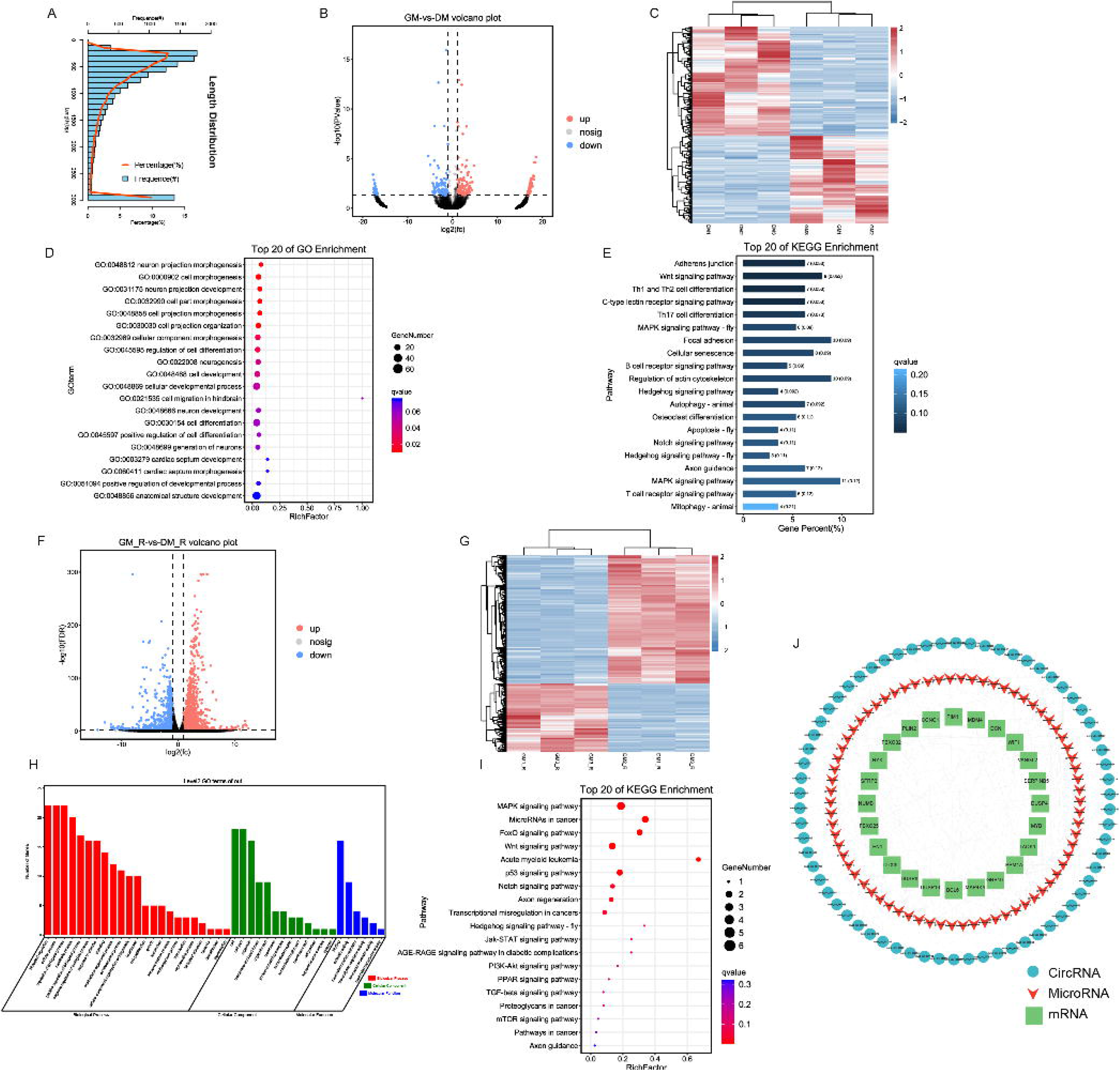
The circRNA-miRNA-mRNA interaction network was constructed by circular RNA sequencing and ribosomal footprinting. (A) circRNA length distribution in GM relative to DM. (B-C) Volcano plot (B) and heatmap (C) of differentially expressed circRNA between proliferation relative to differentiation in chicken myoblasts. (D-E). GO functions (D) and KEGG pathways (E) analysis of the parental genes of differentially expressed circRNAs. (F-G) Volcano plot (F) and heatmap (G) of differentially expressed mRNAs between GM in relative to DM in chicken myoblasts. (H-I) GO functions (F) and KEGG pathways (I) analysis of differentially expressed mRNAs. (J) circRNA-miRNA-mRNA regulatory axes network associating with muscle development, with blue circles representing circRNAs, red triangles representing miRNAs, and green rectangles representing mRNAs.

Combined with the newly identified circRNA and mRNA with differentiated abundance, we characterized a circRNA-miRNA-mRNA interaction network that may involve in regulating the myogenesis process. To be specific, 314 circRNA-miRNA-mRNA regulatory networks with 66 circRNAs, 70 miRNAs, and 24 mRNAs which may be related to myogenesis were acquired as candidate regulation network for further study (Additional file 1).

### *CircPLXNA2* is a novel-identified circRNA regulated by *PLXNA2*

The circPLXNA2-gga-miR-12207-5P-MDM4 axis was the one that piqued our interest for research purposes among these interaction axes. The *circPLXNA2* was termed on the basis of its transcriptional precursor (i.e., pre-PLXNA2-mRNA). Sanger sequencing was initially used to validate the *circPLXNA2* backsplicing junction (Fig 2A). We then validated the circular structure of *circPLXNA2*. We designed specific divergent and convergent primers and used genomic DNA (gDNA) and cDNA as templates respectively for PCR reaction to verify the genomic structure. The expected double bands were inclusively shown in the cDNA group visualized by agarose gel electrophoresis, while when we selected gDNA as an amplifying template, the expected band merely was merely shown in the convergent primers’ lane with weakly detection (Fig 2B). In addition, we employed RNase R treatment to detect the resistance of *circPLXNA2*, which showed successful amplification of circular transcripts by divergent primers using the RNase R-treated cDNA as a template, whereas it is failed to be amplified by using convergent primers. In contrast, targeted bands were successfully amplified by both primers using the cDNA template without RNase R digestion (Figure 2C). We also quantified the RNA expression of the *circPLXNA2* after RNase R treatment, which showed no significant effect on the expression level of *circPLXNA2* after digestion, whereas the expression of β-actin was significantly decreased after digestion (Fig 2C). Together, both findings supported *circPLXNA2’s* actual existence as a circular structure. Nuclear and cytoplasmic localization experiments of *circPLXNA2* were performed to further identify its distribution, and the results showed that *circPLXNA2* was present in both the nucleus and the cytoplasm, with its primary localization being in the latter. (Figure 2F).

**Fig.2.**
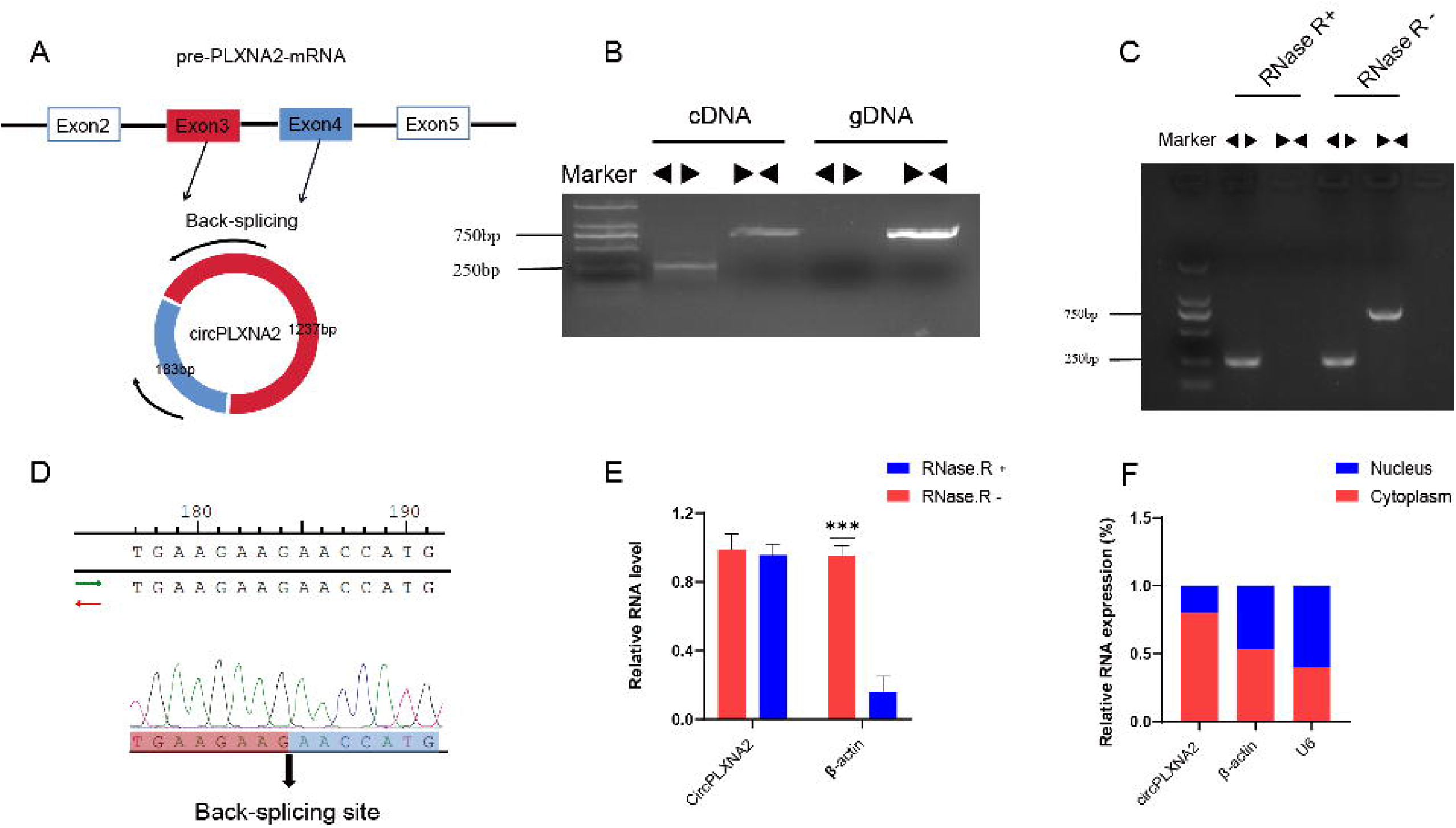
Identification of *circPLXNA2*. (A) Schematic diagram of the cyclization of *pre-PLXNA2-mRNA*. (B) The divergent primers could only amplify *circPLXNA2* in cDNA but not in gDNA visualized in the agarose gel electrophoresis. (C) Convergent primers failed to amplify the target band with RNase R-treated cDNA, whereas divergent primers could successfully amplify the target bands visualized in the agarose gel electrophoresis. (D) Sanger sequencing confirmed the back-splicing junction sequence of *circPLXNA2*. (E) Relative mRNA expression of *circPLXNA2* and β-actin after RNase R treatment. (F) The location of *circPLXNA2* in the cytoplasm and nucleus of CPM was analyzed by qRT-PCR, and GAPDH and U6 were served as cytoplasmic and nuclear localization controls, respectively. (* p < 0.05; ** p < 0.01, *** p < 0.001).

### *CircPLXNA2* promotes cell proliferation and inhibits cell apoptosis

To identify the potential biological function of the *circPLXNA2* in myogenesis, we constructed the overexpression vector of *circPLXNA2*. As shown in Figure 3A, the relative mRNA expression level of *circPLXNA2* was drastically increased after transfecting *PCD25 circPLXNA2* (Fig 3A). Overexpression of *circPLXNA2* in chicken primary myoblasts (CPMs) increased the relative expression of proliferation biomarker genes (*cyclin D1*, *cyclin D2*, and *cyclin B2*), suggesting that it may contribute to proliferation (Fig 3B). Furthermore, we detected the relative expression level of apoptosis-related genes, which revealed that overexpression of *circPLXNA2* suppressed the relative expression of *caspase 3*, *caspase 8*, and *caspase 9* at the mRNA level, indicating detectable apoptosis suppression by *circPLXNA2* (Fig 3C,F). In addition, more 5-ethynyl-2’-deoxyuridine (EdU)-stained cells were found in the *circPLXNA2* overexpression group than in the control group (*PDC25 NC*) (Fig 3D-E). These findings together suggested that *circPLXNA2* could control the development of CPMs by regulating cell proliferation and apoptosis. As we predicted an interaction network involving miRNA (i.e., *gga-miR-12207-5P*), we hypothesized that *circPLXNA2* could serve as an endogenous sponge to competitively interact with *gga-miR-12207-5P*, thereby altering the distribution of *gga-miR-12207-5P* against their target. We used the dual-luciferase reporter assay and the pull-down assay to characterize their potential interaction. Compared with *gga-miR-12207-5P mimic* and *Pmir-GLO-circPLXNA2-MUT* co-transfection group, *gga-miR-12207-5P mimic* co-transfected with *Pmir-GLO-circPLXNA2-WT* vector significantly repressed the relative luciferase activity (Fig 3H). Gel electrophoresis and biotin-labeled miRNA pull-down analysis demonstrated that *biotin-gga-miR-12207-5P mimic* transfection significantly decreased the *circPLXNA2* mRNA level in CPMs (Fig 3I-J). Collectively, these findings suggested that *circPLXNA2* and *gga-miR-12207-5P* have a targeted binding relationship.

**Fig.3.**
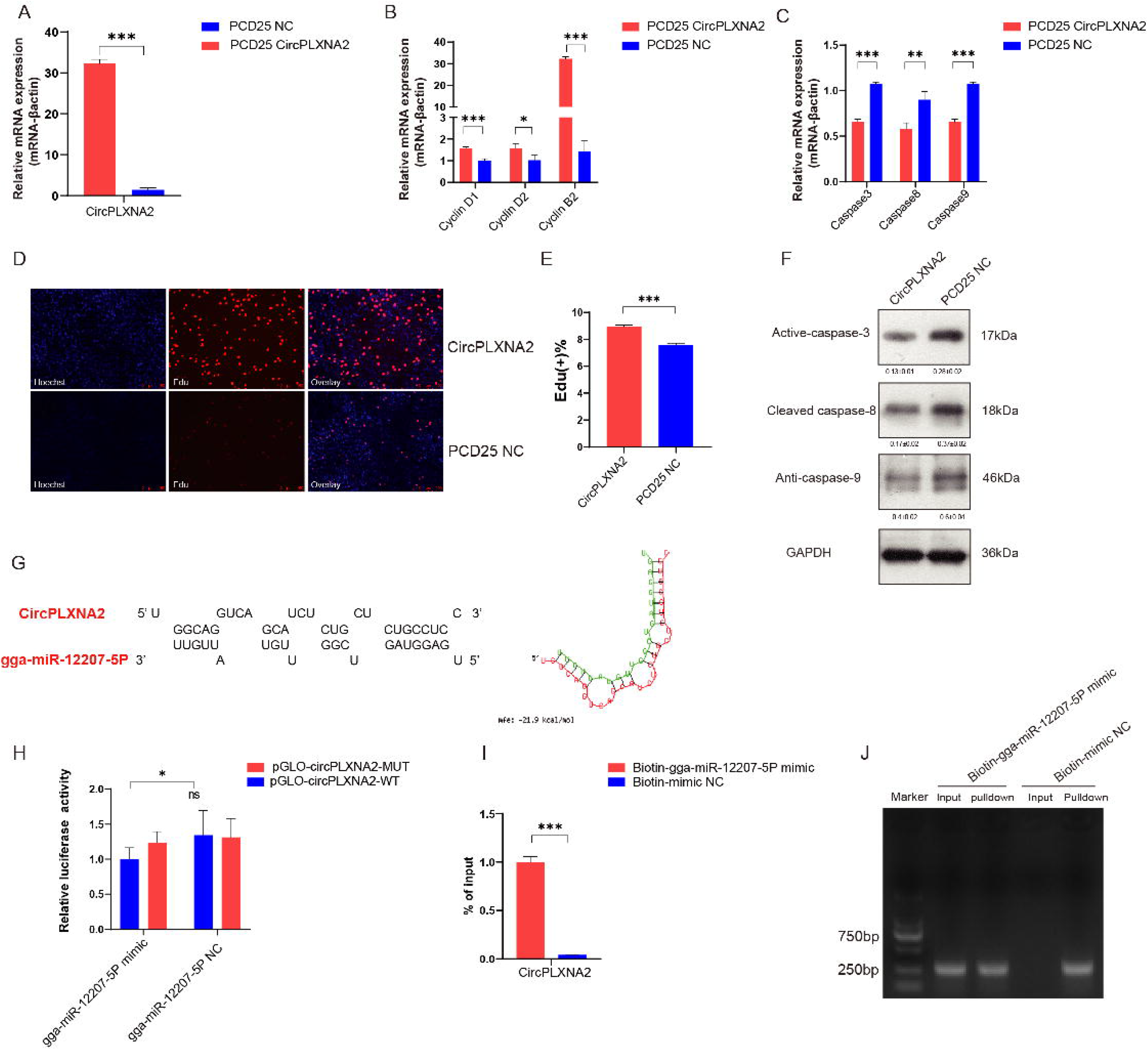
*CircPLXNA2* promoted cell proliferation, inhibited cell apoptosis and sponged *gga-miR-12207-5P*. (A) Efficiency of overexpression of *circPLXNA2*. (B) qRT-PCR results of *Cyclin B2*, *cyclin D1*, *cyclin D2*, and *cyclin B2* genes after *circPLXNA2* overexpression. (C) qRT-PCR results of *caspase 3, caspase 8* and *caspase 9* genes after *circPLXNA2* overexpression. (D-E) EdU staining (D) and the positive EdU cell rate (E) for myoblast cell after *circPLXNA2* overexpression. (F) Western blotting analysis of the *caspase 3, caspase 8*, and *caspase 9* proteins with PCD25 *circPLXNA2* transfection. (G) Potential binding sites between *circPLXNA2* and *gga-miR-12207-5P* predicted by RNAhybrid online tool. (H) Relative luciferase activity of *pmirGLO-circPLXNA2* WT/MUT plasmid with *gga-miR-12207-5P mimic* or *gga-miR-12207-5P* NC. (I-J) The biotin-tagged miRNA pulldown showed the interaction between *circPLXNA2* and *gga-miR-12207-5P*. (* p < 0.05; ** p < 0.01, *** p < 0.001).

### *gga-miR-12207-5P* inhibits cell proliferation and promotes apoptosis

To explore the biological function of *gga-miR-12207-5P*, we designed the overexpression vector of *gga-miR-12207-5P* and termed *gga-miR-12207-5P-mimic*. After 48h of transfection, the relative expression level of *gga-miR-12207-5P* was significantly higher than the control group, indicating reliable overexpression efficiency. (Fig 4A). After transferring *gga-miR-12207-5P-mimic* into CPMs 48h, *cyclin D1*, *cyclin D2*, and *cyclin B2* were all significantly downregulated compared to *mimic NC* treatment, suggesting that it inhibits proliferation (Fig 4B). After 48 hours of transfection, the apoptosis pathway was activated, as evidenced by an increase in the relative expression level of *caspase 8*, *caspase 9*, and *caspase 3* (Fig 4C-D). Edu staining further backbone the fact that *gga-miR-12207-5P* overexpression significantly repressed the viability of CPMs (Fig 4E-F). Overall, the viability of CPMs may be downregulated by overexpression *gga-miR-12207-5P*.

**Fig.4.**
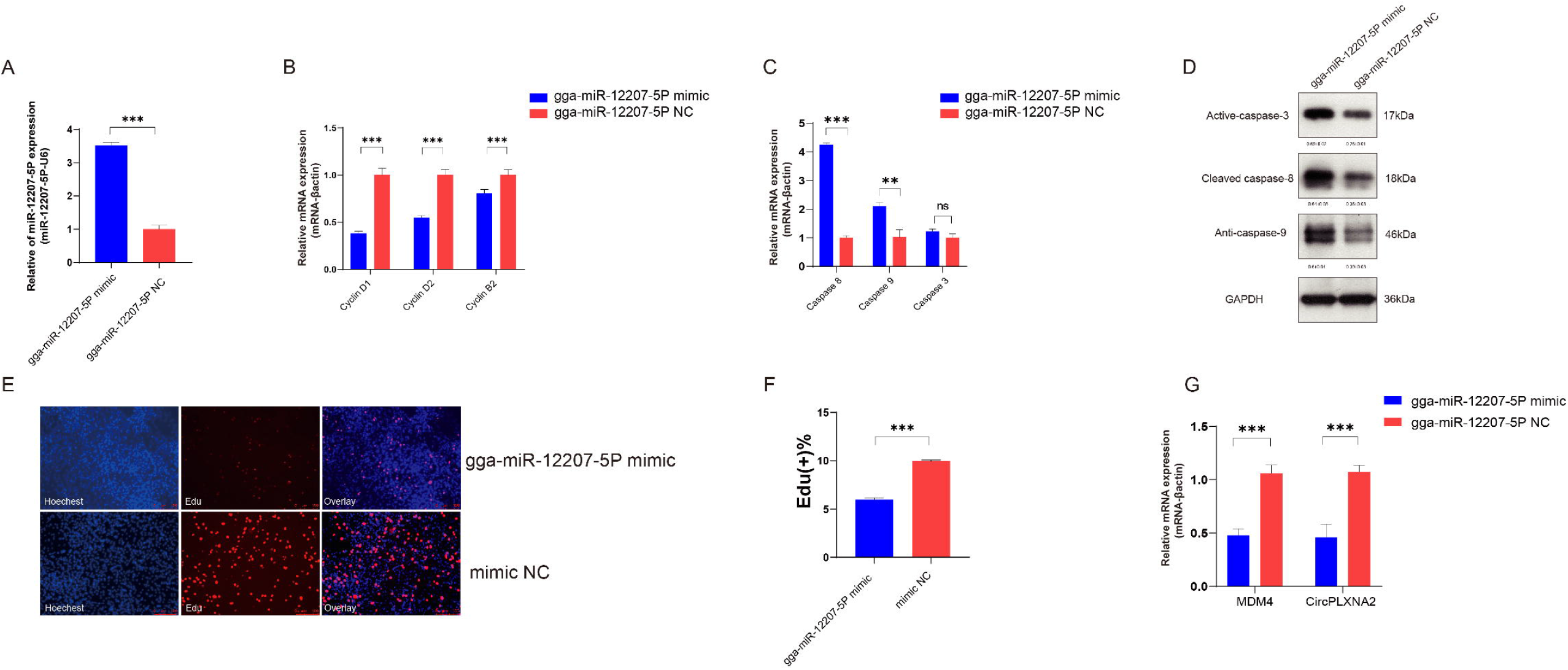
*gga-miR-12207-5P* inhibited cell proliferation and promoted cell apoptosis. (A) The overexpression efficiency of *gga-miR-12207-5P* in myoblasts after transfection with *gga-miR-12207-5P mimic*. (B) qRT-PCR results of *cyclin D1, cyclin D2*, and *cyclin B2* after *gga-miR-12207-5P* overexpression. (C-D) mRNA level (C) and translation level (D) of *Caspase 3, Caspase 8* and *Caspase 9* after *gga-miR-12207-5P* overexpression. (E-F) EdU staining (E) and the positive EdU cell rate (F) for myoblast cell after *gga-miR-12207-5P* overexpression. (G) The expression levels of *circPLXNA2* and *MDM4* were significantly decreased in myoblasts transfected with *gga-miR-12207-5P mimic*. (* p < 0.05; ** p < 0.01, *** p < 0.001).

Additionally, we verified the potential interaction between *gga-miR-12207-5P, MDM4*, and *circPLXNA2*. Overexpression of *gga-miR-12207-5P* decreased *MDM4* and *circPLXNA2* mRNA abundance, suggesting that it inhibits their transcriptional activity (Fig 4G).

### *MDM4* promotes cell proliferation and inhibits cell apoptosis through the *P53* signaling pathway

To confirm *MDM4*’s biological function in the myogenesis process, we created an overexpression plasmid of *MDM4* to detect and transfected into CPMs, which showed significant repression of relative mRNA expression of *p53*, a critical signaling pathway regulating cell apoptosis, indicating that *MDM4* may inhibit myogenesis via apoptosis. (Fig 5A). As such, we detected the relative expression level of *cyclin D1*, *cyclin D2*, and *cyclin D3* when *MDM4* was overexpressed in CPMs. The results revealed that these genes were significantly promoting expression at the mRNA level. EdU staining assays also suggested that *MDM4* overexpression induced CPM proliferation (Fig 5B-D). On the contrary, silencing the *MDM4* gene repressed the relative mRNA expression of *cyclin D1*, *cyclin D2*, and *cyclin D3*, as well as the total number of stained cells (Fig 5E-H).Furthermore, we detected the relative expression level of apoptosis-related genes and found that *caspase 3*, *caspase 8*, and *caspase 9* were significantly downregulated after 48h transfecting an overexpression *MDM4* plasmid (I). Western blotting analysis also showed a decreasing trend of *caspase 3*, *caspase 8*, and *caspase 9* after *MDM4* overexpression (J). Conversely, we observed opposite effect of *caspase 3*, *caspase 8*, and *caspase 9* as the expression of *MDM4* was interfered (Fig 5K). Congruously, these results supported the biological role of *MDM4* in promoting cell proliferation and inhibiting apoptosis. We used the RNA hybrid tool to predict the target binding region of 3’UTR of *MDM4* and *gga-miR-12207-5P* seed sequence and identified the best-matched region possessing the most minimum free energy (Fig 5L). Subsequently, dual-luciferase reporter vectors (i.e., Pmir-*GLO-MDM4-W*T and *Pmir-GLO-MDM4-MUT*) were constructed and transferred into DF-1 cells to further explore the potential interaction of *MDM4* and *gga-miR-12207-5P*. RT-qPCR results demonstrated the targeted binding relationship between *gga-miR-12207-5P* and *MDM4* (Fig 5M), which was further confirmed by a biotin-coupled miRNA pull-down assay (Fig 5N-O).

**Fig.5.**
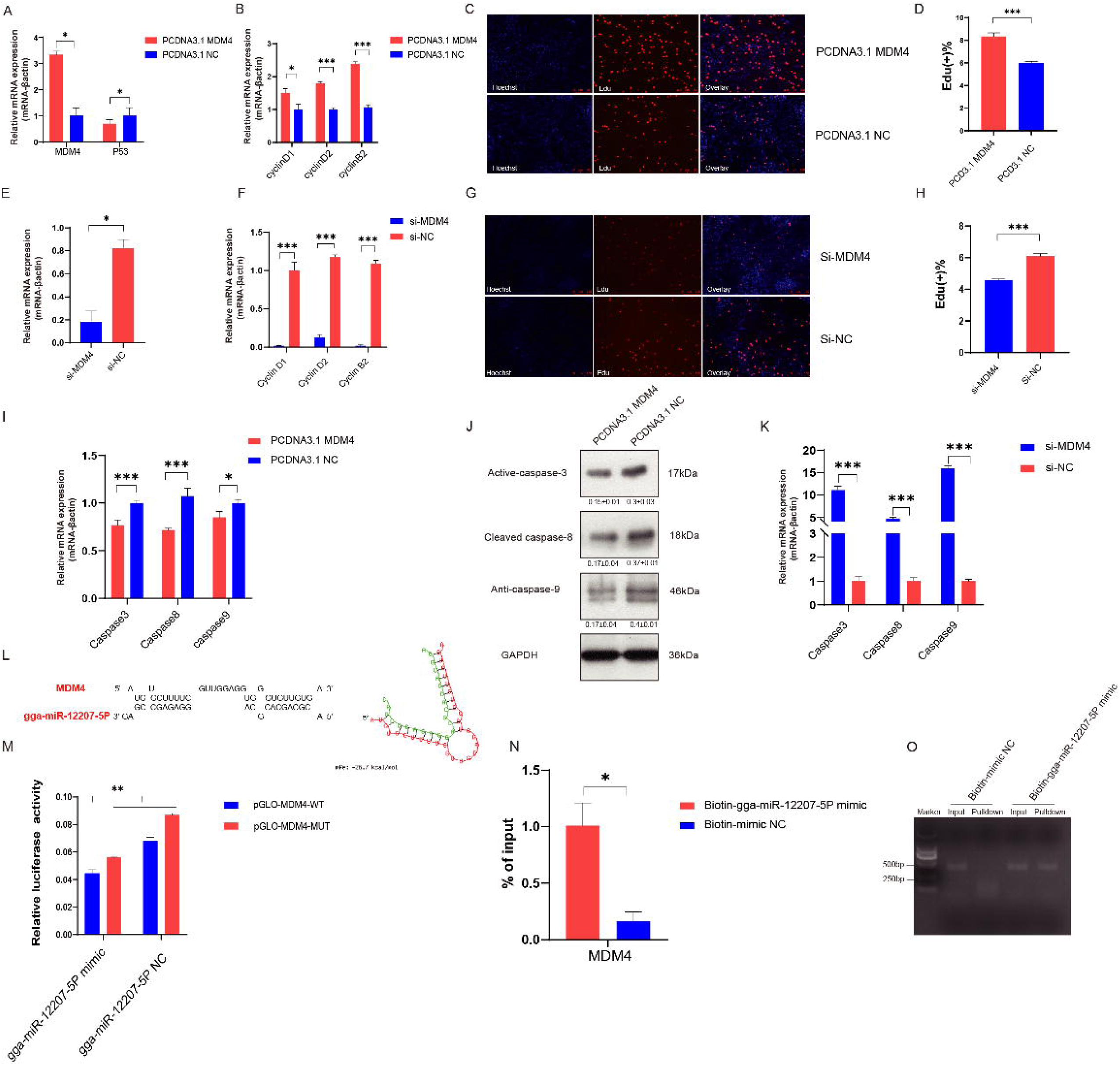
*MDM4* promotes cell proliferation and inhibits apoptosis through *P53* signaling pathway. (A) *MDM4* and *p53* mRNA levels after transfection with *MDM4* overexpression plasmid in CPMs. (B) qRT-PCR results of *cyclin D1*, *cyclin D2*, and *cyclin B2* after *MDM4* overexpression. (C-D) EdU staining (C) and the positive EdU cell rate (D) for myoblast cell after *MDM4* overexpression. (E) *MDM4* mRNA levels after transfection with *MDM4* silencing vector in CPMs. (F) qRT-PCR results of *cyclin D1*, *cyclin D2*, and *cyclin B2* after *MDM4* silencing. (G-H) EdU staining (G) and the positive EdU cell rate (H) for myoblast cell after *MDM4* knockdown. (I-J) relative mRNA level (I) and protein level (J) of *caspase 3, caspase 8*, and *caspase 9* expression after *MDM4*overexpression. (K) relative mRNA level expression of *caspase 3*, *caspase 8*, and *caspase 9* genes after *MDM4* gene silence. (L) Potential binding sites between *MDM4* and *gga-miR-12207-5P* predicted by RNAhybrid online tool. (M) Relative luciferase activity of *pmirGLO-MDM4 WT/MUT* plasmid with *gga-miR-12207-5P mimic* or *gga-miR-12207-5P* NC. (N-O) mRNA level (N) and RNA level (O) of the biotin-tagged miRNA pulldown assay. (* p < 0.05; ** p < 0.01, *** p < 0.001).

### *CircPLXNA2* affects the proliferation and apoptosis of myoblasts by regulating *MDM4* through competitive sponge of *gga-miR-12207-5P*

To further characterize the potential interaction relationship between *circPLXNA2*, *gga-miR-12207-5P*, and *MDM4*, we measured *MDM4* expression levels after overexpressing both *gga-miR-12207-5P mimic* and *circPLXNA2*. To be more specific, compared to the *gga-miR-12207-5P mimic* and *circPLXNA2* co-transfection groups, the mRNA expression level of *MDM4* was significantly upregulated in the *gga-miR-12207-5P mimic* and *PCD25* co-transfection groups, as well as the *mimic NC* and *circPLXNA2* co-transfection groups. Also, the relative mRNA expression level of *MDM4* was lowest in the group that had both *gga-miR-12207-5P mimic* and PCD25 co-transfected. In contrast, the expression level of *MDM4* was highest when *circPLXNA2* was overexpressed, which suggests that *circPLXNA2* may reverse the effect of gga-miR-12207-5P on MDM4 (Fig 6A).

**Fig.6.**
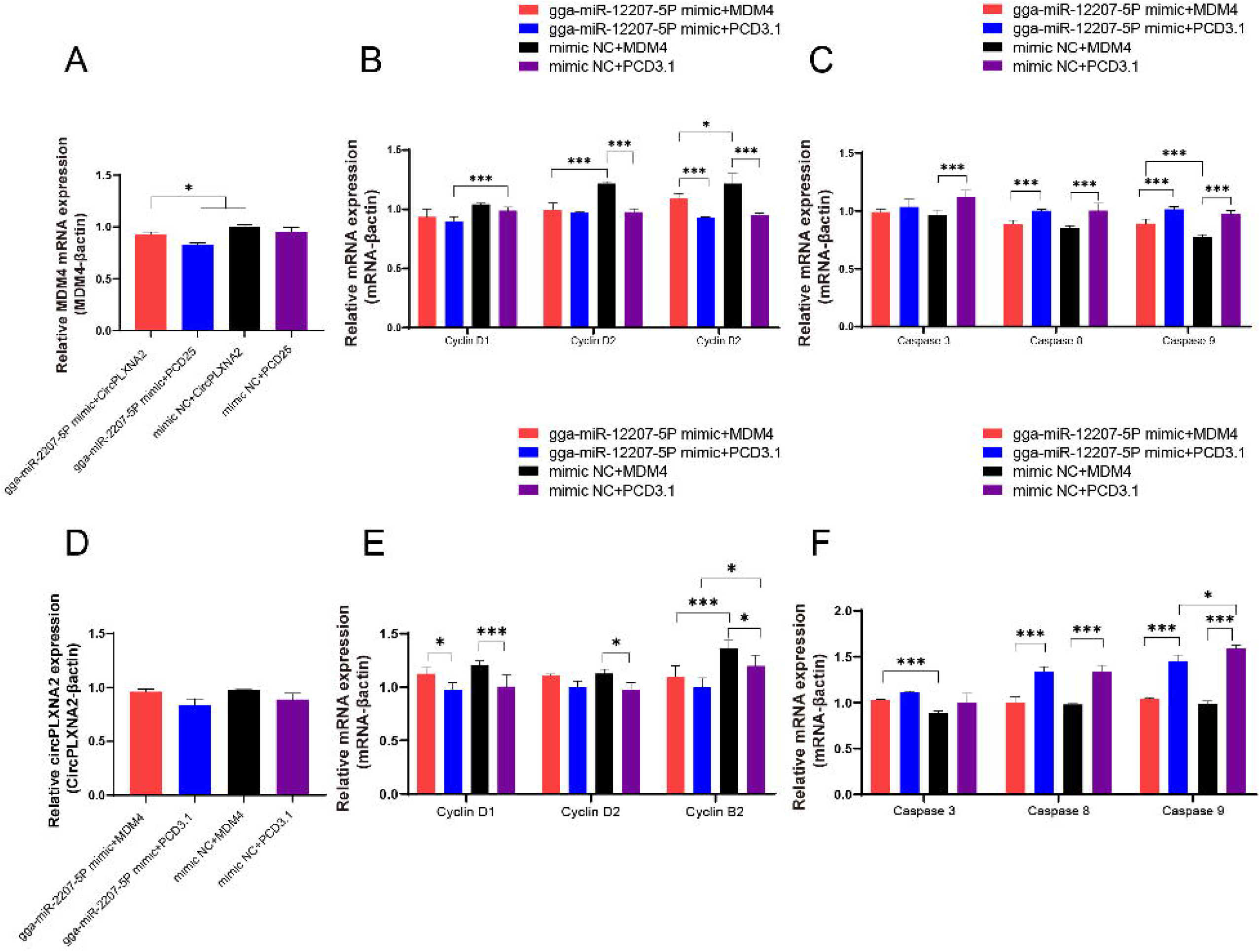
*CircPLXNA2* regulates *MDM4* by competitive adsorption of *gga-miR-12207-5P* to affect the proliferation and apoptosis of myoblasts. (A) The mRNA expression level of *MDM4* after co-transfection of *gga-miR-12207-5P mimic* and *circPLXNA2* in myoblasts. (B) The mRNA expression levels of *cyclin D1*, *cyclin D2*, and *cyclin B2* after co-transfection of *gga-miR-12207-5P mimic* and *circPLXNA2* in myoblasts. (C) The mRNA expression levels of *caspase 3*, *caspase 8*, and *caspase 9* after co-transfection of *gga-miR-12207-5P mimic* and *circPLXNA2* in myoblasts. (D) Expression level of *circPLXNA2* after co-transfection of *gga-miR-12207-5P mimic* and *MDM4* in myoblasts. (E) qRT-PCR results of *cyclin D1*, *cyclin D2* and, *cyclin B2* after *gga-miR-12207-5P mimic* and *MDM4* co-transfection. (F) qRT-PCR results of *caspase 3*, *caspase 8*, and *caspase 9* after *gga-miR-12207-5P mimic* and *MDM4* co-transfection.

**Fig.7.**
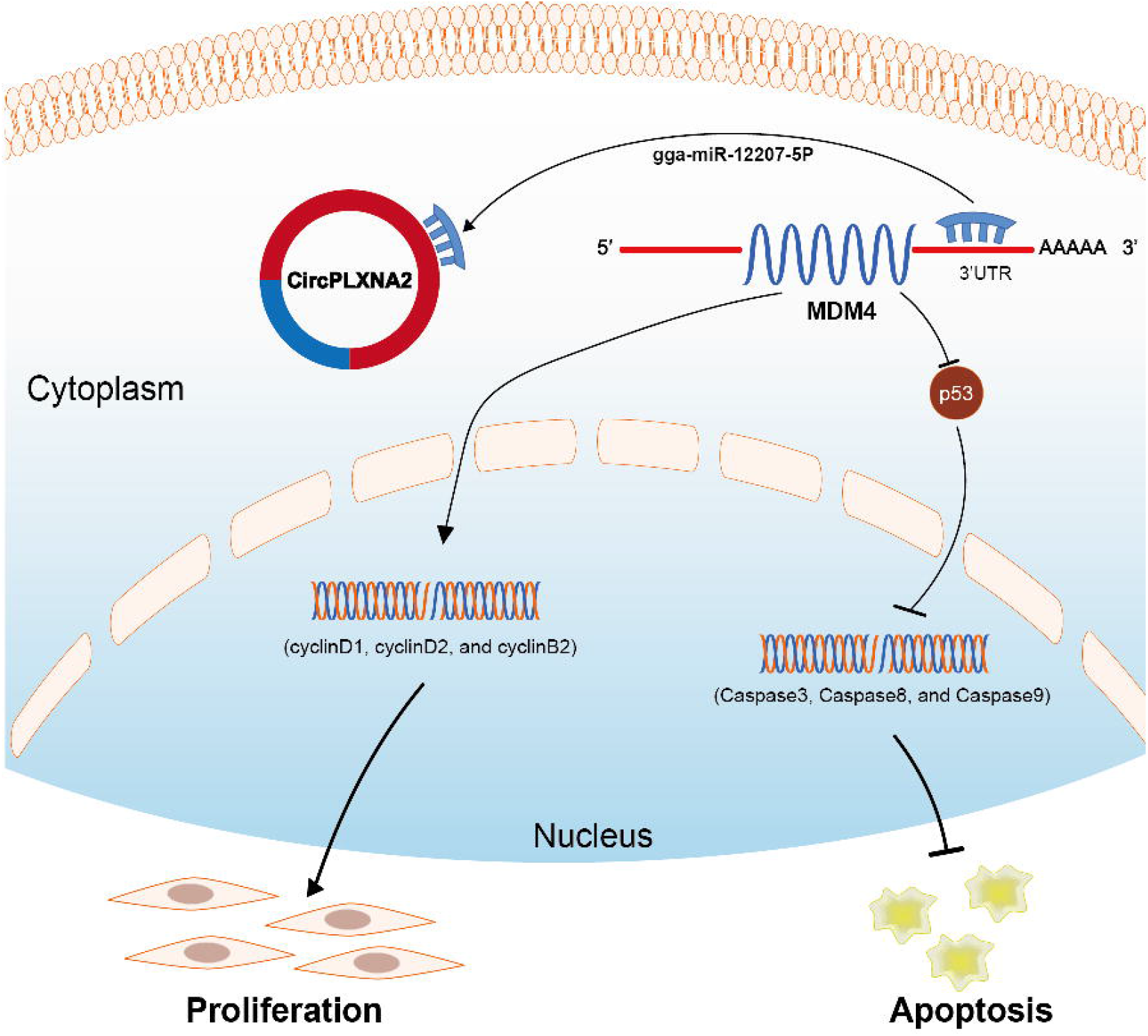
Schematic diagram of *CircPLXNA2* affected the proliferation and apoptosis of myoblast *circPLXNA2/gga-miR-12207-5P/MDM4* axis.

We further quantified the *cyclin B2*, *cyclin D1*, and *cyclin D2* genes after overexpressing *gga-miR-12207-5P* and *circPLXNA2* in CPMs, individually or in combination, to confirm the potential proliferation effect of both molecules. As shown, the RT-qPCR results indicated that as *circPLXNA2* was overexpressed independently, the mRNA relative expression of the three genes was drastically higher than other groups (*p<0.05*), whereas *gga-miR-12207-5P* independent overexpression reduced the mRNA relative expression of the three genes. We therefore co-transfected *circPLXNA2* and *gga-miR-12207-5P mimic*, and discovered that the activity levels of these three genes were partially restored (Fig 6B). As well, we measured the abundance of apoptosis markers *caspase 3*, *caspase 8*, and *caspase 9* mRNAs after overexpression *gga-miR-12207-5P* and *circPLXNA2* respectively, or co-overexpression of both. RT-qPCR measurements showed that when *circPLXNA2* was overexpressed alone, the three apoptosis markers were drastically downregulated compared to the other three groups. However, when we transfected the *gga-miR-12207-5P* mimic alone, the mRNA abundance of the three markers increased significantly (p<0.05). Subsequently, we co-transfected both *gga-miR-12207-5P mimic* and *circPLXNA2*, and *caspase 3*, *caspase 8*, and *caspase 9* were reverted between transfections. (Fig 6C). These findings led us to hypothesize that *circPLXNA2* could free *MDM4* by competing with it for binding to *gga-miR-12207-5P*, restoring *MDM4*’s ability to promote cell proliferation while simultaneously inhibiting apoptosis. In order to measure the levels of *circPLXNA2*, we co-transfected *gga-miR-12207-5P mimic* and *MDM4*. We discovered a similar tendency when compared to the *MDM4* expression levels as we transfected *gga-miR-12207-5P mimic* and *circPLXNA2* separately or both synchronously. We also measured the relative levels of proliferation and apoptosis markers after *gga-miR-12207-5P* and *circPLXNA2* co-transfection or independent transfection, which showed a close variable trend (Fig 6 B-C). Overall, these results back our hypothesis, that is, *circPLXNA2* recovered the function of *MDM4* by competitively binding to *gga-miR-12207-5P*, thereby recovering the promotion of cell proliferation and inhibition of apoptosis by *MDM4*.

## Discussion

CircRNAs are new members of the non-coding RNA family that emerged from the backbone of pre-mRNA back splicing and were previously classified as “transcriptional noise” in mammals [24, 25]. Furthermore, due to the circRNA’s low abundance and expressive quantity, the conventional mode of study is inapplicable [26]. However, with the progress of the genome sequencing approach, circRNAs are increasingly recognized to be implicated in regulating disease phenotype. Depending on their subcellular location, circRNAs are regulated in two distinct ways. [27]. In the nucleus, circRNAs would combine with RNA polymerase ʡ to regulate the transcriptional activity [28]. Alternatively, in the cytoplasm, circRNAs would act as molecular sponges to bind with specific miRNA to promote the function of target genes [5, 13, 20, 29–34]. For example, it has been demonstrated that *circPTPN4* can stimulate NAMPT expression by sponging *miR-499-3P* in the cytoplasm and acting as a competing endogenous RNA [13]. Hence, here we favor a model to identify the circRNA-miRNA-mRNA interaction network in which the axes may be implicated in the progress of CPMs myogenesis in a combination of multi-omics (i.e., circRNA-seq and ribo-seq). To be more specific, we first used circRNA-seq to identify a total of 360 circRNAs with differential expression during the GM compared to the DM. We further presented the 3342 ribo-seq-identified differentially-transcribed genes, thereby selecting the genes most likely to play a role in myogenesis during the GM as opposed to the DM. Combining these, we characterized a circRNA-miRNA-mRNA network that may participate in regulating myogenesis.

In our study, we identified a novel circRNA with heterogeneous expression in GM relative to DM and termed *circPLXNA2*, which is generated from 3-4 exon of *PLXNA2* gene. Overexpression, real-time quantitative PCR, a dual-luciferase reporter assay, RNA pull-down, and the addition of EdU were the methods that we used to demonstrate the potential biological role of *circPLXNA2*.We found that *circPLXNA2* could promote the relative mRNA abundance of *cyclin D1, cyclin D2*,and *cyclin D3* and inhibit mRNA abundance of *caspase 3, caspase 8*, and *caspase 9*, facilitating cell proliferation and repress apoptosis. In addition, the interaction between gga-miR-12207-5P and circPLXNA2 was investigated further and found to be consistent with the results of our in-silico analysis.

*MDM4* is constituted as a *p53* inhibitor to regulate *p53* activity, thereby inhibiting cell apoptosis and promoting proliferation [20]. *P53* is a sensor of genotoxic stress that could protect cells from DNA damage by inducing cell-cycle arrest [21]. Numerous studies have shown that *p53* induces apoptosis via both transcriptional activation of pro-apoptotic and repression of anti-apoptotic genes, as well as non-transcriptional mechanisms [14, 15]. As exemplified by Cai et al, *miR-16-5p* could inhibit proliferation and promote apoptosis by *p53* signaling pathway via regulating *SESN1* mRNA relative expression [22]. Here, we showed that *gga-miR-12207-5P* has the opposite effect on *circPLXNA2*, namely, that it can suppress *MDM4*, which controls myoblast proliferation and apoptosis via the *p53* signaling pathway. By simultaneously overexpressing *circPLXNA2* and *gga-miR-12207-5P*, we were able to restore the relative abundance of *MDM4*, which in turn reversed proliferation and inhibited apoptosis.

## Conclusion

Using circRNA-seq and in combination with ribo-seq, we analyzed the differentially expressed circRNA in myoblasts during GM relative DM, thereby building an interactive regulation network associated with myoblast development. Through this, a novel regulatory axis known as circPLXNA2/gga-miR-12207-5P/MDM4 and its biological function were further characterized. To be specific, *circPLXNA2* could function as ceRNA to sponge *gga-miR-12207-5P* to release *MDM4*, a negative regulator of *p53* signaling pathway, thereby exerting the myoblast proliferation and repressing apoptosis. Overall, our study provided a comprehensive profiling of interaction axes which may regulate in the process of myogenesis. As well, this study showed the first evidence of *circPLXNA2* of regulating skeletal myogenesis, and the circPLXNA2-gga-miR-12207-5p-MDM4 may become a potential therapeutic target for muscle development.

## Acknowledgments

This work was supported by the National Key R & D Program of China (Grant No. 2021YFD1300100) and Science and Technology Program of Guangzhou, China (Grant No. 202201010507).

## Author Contributions

Conceptualization, methodology, Zhenhui Li, Xu Dong; software, Xu Dong; validation, Xu Dong, Jiabao Xing; formal analysis, Zhen Zhou; data curation, original draft preparation, Xu Dong, Jiabao Xing; revision and editing, Qinghua Nie, Zhenhui Li; visualization, Xu Dong, Jiabao Xing; supervision, Qinghua Nie, Zhuihui Li; funding acquisition, Qinghua Nie, Zhuihui Li. All authors have read and agreed to the published version of the manuscript.

## Conflicts of Interest

The authors declare no conflict of interest.

## Availability of data and materials

The circRNA-seq and the ribo-seq profiles have been deposited in the China National GeneBank DataBase (CNGBdb) under the accession number CNP0003882. Any remaining data could be acquired in the supplement files or requested to the corresponding author.

## Abbreviations

CeRNA: Competing endogenous RNA
circRNA: Circular RNA
DM: Differentiation medium
CPMs: Chicken primary myoblasts
EdU: 5-Ethynyl-2’-deoxyuridine;
gDNA: Genomic DNA
GO: Gene Ontology
IF: Immunofluorescence
KEGG: Kyoto Encyclopedia of Genes and Genomes
miRNA: MicroRNA
NC: Negative control
ncRNA: Noncoding RNA
siRNA: Smal interfering RNA
UTR: Untranslated region
MDM4: Murine Double Minute 4
DM: Differentiation medium
GM: Growth medium

## Additional file

Additional file 1 Table showing the detail of identified interaction network by using multi-omics.

Additional file 2 Chromosome distribution of circRNA transcripts identified in GM relative to DM of chicken primary myogenesis.

Additional file 3 Backspliced reads distribution of circRNA transcripts identified in GM relative to DM of chicken primary myogenesis.

Additional file 4 Differentiated-expressed circRNAs identified in GM relative to DM of chicken primary myogenesis by using circRNA-seq.

Additional file 5 Differentiated-expressed genes identified in GM relative to DM of chicken primary myogenesis by using ribo-seq.

## Notes

### Competing Interest Statement

The authors have declared no competing interest.

## Reference

1. Güller I, Russell AP: MicroRNAs in skeletal muscle: their role and regulation in development, disease and function. J Physiol 2010, 588(Pt 21):4075–4087.

2. Das A, Das A, Das D, Abdelmohsen K, Panda AC: Circular RNAs in myogenesis. Biochim Biophys Acta Gene Regul Mech 2020, 1863(4):194372.

3. Zhang P, Chao Z, Zhang R, Ding R, Wang Y, Wu W, Han Q, Li C, Xu H, Wang L et al: Circular RNA Regulation of Myogenesis. Cells 2019, 8(8).

4. Piwecka M, Glažar P, Hernandez-Miranda LR, Memczak S, Wolf SA, Rybak-Wolf A, Filipchyk A, Klironomos F, Cerda Jara CA, Fenske P et al: Loss of a mammalian circular RNA locus causes miRNA deregulation and affects brain function. Science 2017, 357(6357).

5. Yan J, Yang Y, Fan X, Liang G, Wang Z, Li J, Wang L, Chen Y, Adetula AA, Tang Y et al: circRNAome profiling reveals circFgfr2 regulates myogenesis and muscle regeneration via a feedback loop. J Cachexia Sarcopenia Muscle 2022, 13(1):696–712.

6. Hansen TB, Jensen TI, Clausen BH, Bramsen JB, Finsen B, Damgaard CK, Kjems J: Natural RNA circles function as efficient microRNA sponges. Nature 2013, 495(7441):384–388.

7. Memczak S, Jens M, Elefsinioti A, Torti F, Krueger J, Rybak A, Maier L, Mackowiak SD, Gregersen LH, Munschauer M et al: Circular RNAs are a large class of animal RNAs with regulatory potency. Nature 2013, 495(7441):333–338.

8. Wang Y, Li M, Wang Y, Liu J, Zhang M, Fang X, Chen H, Zhang C: A Zfp609 circular RNA regulates myoblast differentiation by sponging miR-194-5p. Int J Biol Macromol 2019, 121:1308–1313.

9. Wei X, Li H, Yang J, Hao D, Dong D, Huang Y, Lan X, Plath M, Lei C, Lin F et al: Circular RNA profiling reveals an abundant circLMO7 that regulates myoblasts differentiation and survival by sponging miR-378a-3p. Cell Death Dis 2017, 8(10):e3153.

10. Lee RC, Feinbaum RL, Ambros V: The C. elegans heterochronic gene lin-4 encodes small RNAs with antisense complementarity to lin-14. Cell 1993, 75(5):843–854.

11. Yang X, Li Z, Wang Z, Yu J, Ma M, Nie Q: miR-27b-3p Attenuates Muscle Atrophy by Targeting Cbl-b in Skeletal Muscles. Biomolecules 2022, 12(2).

12. Chen JF, Mandel EM, Thomson JM, Wu Q, Callis TE, Hammond SM, Conlon FL, Wang DZ: The role of microRNA-1 and microRNA-133 in skeletal muscle proliferation and differentiation. Nat Genet 2006, 38(2):228–233.

13. Cai B, Ma M, Zhou Z, Kong S, Zhang J, Zhang X, Nie Q: circPTPN4 regulates myogenesis via the miR-499-3p/NAMPT axis. J Anim Sci Biotechnol 2022, 13(1):2.

14. Mercer J, Mahmoudi M, Bennett M: DNA damage, p53, apoptosis and vascular disease. Mutat Res 2007, 621(1-2):75–86.

15. Vousden KH, Lu X: Live or let die: the cell’s response to p53. Nat Rev Cancer 2002, 2(8):594–604.

16. Levine AJ: p53, the cellular gatekeeper for growth and division. Cell 1997, 88(3):323–331.

17. Ashcroft M, Vousden KH: Regulation of p53 stability. Oncogene 1999, 18(53):7637–7643.

18. Oren M: Regulation of the p53 tumor suppressor protein. J Biol Chem 1999, 274(51):36031–36034.

19. Mendrysa SM, McElwee MK, Michalowski J, O’Leary KA, Young KM, Perry ME: mdm2 Is critical for inhibition of p53 during lymphopoiesis and the response to ionizing irradiation. Mol Cell Biol 2003, 23(2):462–472.

20. Mancini F, Gentiletti F, D’Angelo M, Giglio S, Nanni S, D’Angelo C, Farsetti A, Citro G, Sacchi A, Pontecorvi A et al: MDM4 (MDMX) overexpression enhances stabilization of stress-induced p53 and promotes apoptosis. J Biol Chem 2004, 279(9):8169–8180.

21. Hay N: p53 strikes mTORC1 by employing sestrins. Cell Metab 2008, 8(3):184–185.

22. Cai B, Ma M, Chen B, Li Z, Abdalla BA, Nie Q, Zhang X: MiR-16-5p targets SESN1 to regulate the p53 signaling pathway, affecting myoblast proliferation and apoptosis, and is involved in myoblast differentiation. Cell Death Dis 2018, 9(3):367.

23. Li Z, Cai B, Abdalla BA, Zhu X, Zheng M, Han P, Nie Q, Zhang X: LncIRS1 controls muscle atrophy via sponging miR-15 family to activate IGF1-PI3K/AKT pathway. J Cachexia Sarcopenia Muscle 2019, 10(2):391–410.

24. Meng S, Zhou H, Feng Z, Xu Z, Tang Y, Li P, Wu M: CircRNA: functions and properties of a novel potential biomarker for cancer. Mol Cancer 2017, 16(1):94.

25. Xu S, Zhou L, Ponnusamy M, Zhang L, Dong Y, Zhang Y, Wang Q, Liu J, Wang K: A comprehensive review of circRNA: from purification and identification to disease marker potential. PeerJ 2018, 6:e5503.

26. Cocquerelle C, Mascrez B, Hétuin D, Bailleul B: Mis-splicing yields circular RNA molecules. Faseb j 1993, 7(1):155–160.

27. Ebbesen KK, Hansen TB, Kjems J: Insights into circular RNA biology. RNA Biol 2017, 14(8):1035–1045.

28. Salzman J, Gawad C, Wang PL, Lacayo N, Brown PO: Circular RNAs are the predominant transcript isoform from hundreds of human genes in diverse cell types. PLoS One 2012, 7(2):e30733.

29. Wang PL, Bao Y, Yee MC, Barrett SP, Hogan GJ, Olsen MN, Dinneny JR, Brown PO, Salzman J: Circular RNA is expressed across the eukaryotic tree of life. PLoS One 2014, 9(6):e90859.

30. Yue B, Wang J, Ru W, Wu J, Cao X, Yang H, Huang Y, Lan X, Lei C, Huang B et al: The Circular RNA circHUWE1 Sponges the miR-29b-AKT3 Axis to Regulate Myoblast Development. Mol Ther Nucleic Acids 2020, 19:1086–1097.

31. Peng S, Song C, Li H, Cao X, Ma Y, Wang X, Huang Y, Lan X, Lei C, Chaogetu B et al: Circular RNA SNX29 Sponges miR-744 to Regulate Proliferation and Differentiation of Myoblasts by Activating the Wnt5a/Ca(2+) Signaling Pathway. Mol Ther Nucleic Acids 2019, 16:481–493.

32. Chen M, Wei X, Song M, Jiang R, Huang K, Deng Y, Liu Q, Shi D, Li H: Circular RNA circMYBPC1 promotes skeletal muscle differentiation by targeting MyHC. Mol Ther Nucleic Acids 2021, 24:352–368.

33. Huang K, Chen M, Zhong D, Luo X, Feng T, Song M, Chen Y, Wei X, Shi D, Liu Q et al: Circular RNA Profiling Reveals an Abundant circEch1 That Promotes Myogenesis and Differentiation of Bovine Skeletal Muscle. J Agric Food Chem 2021,69(1):592–601.

34. Chen Y, Wang Z, Chen X, Peng X, Nie Q: CircNFIC Balances Inflammation and Apoptosis by Sponging miR-30e-3p and Regulating DENND1B Expression. Genes (Basel) 2021, 12(11).

