## Supplementary figures and images for "*CircPLXNA2* affects the proliferation and apoptosis of myoblast through *circPLXNA2/gga-miR-12207-5P/MDM4* axis"

### Additional file 2

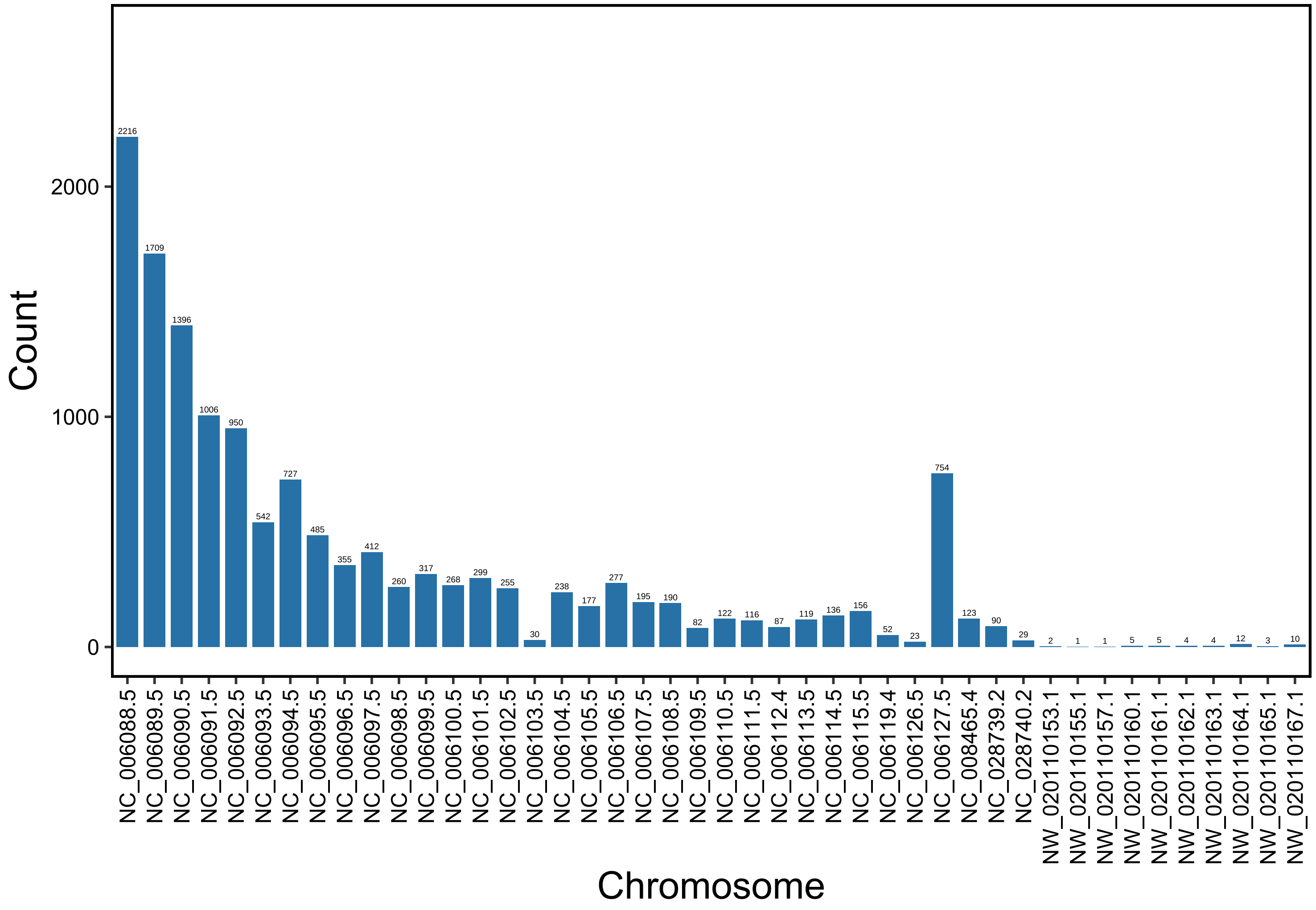

### Additional file 3

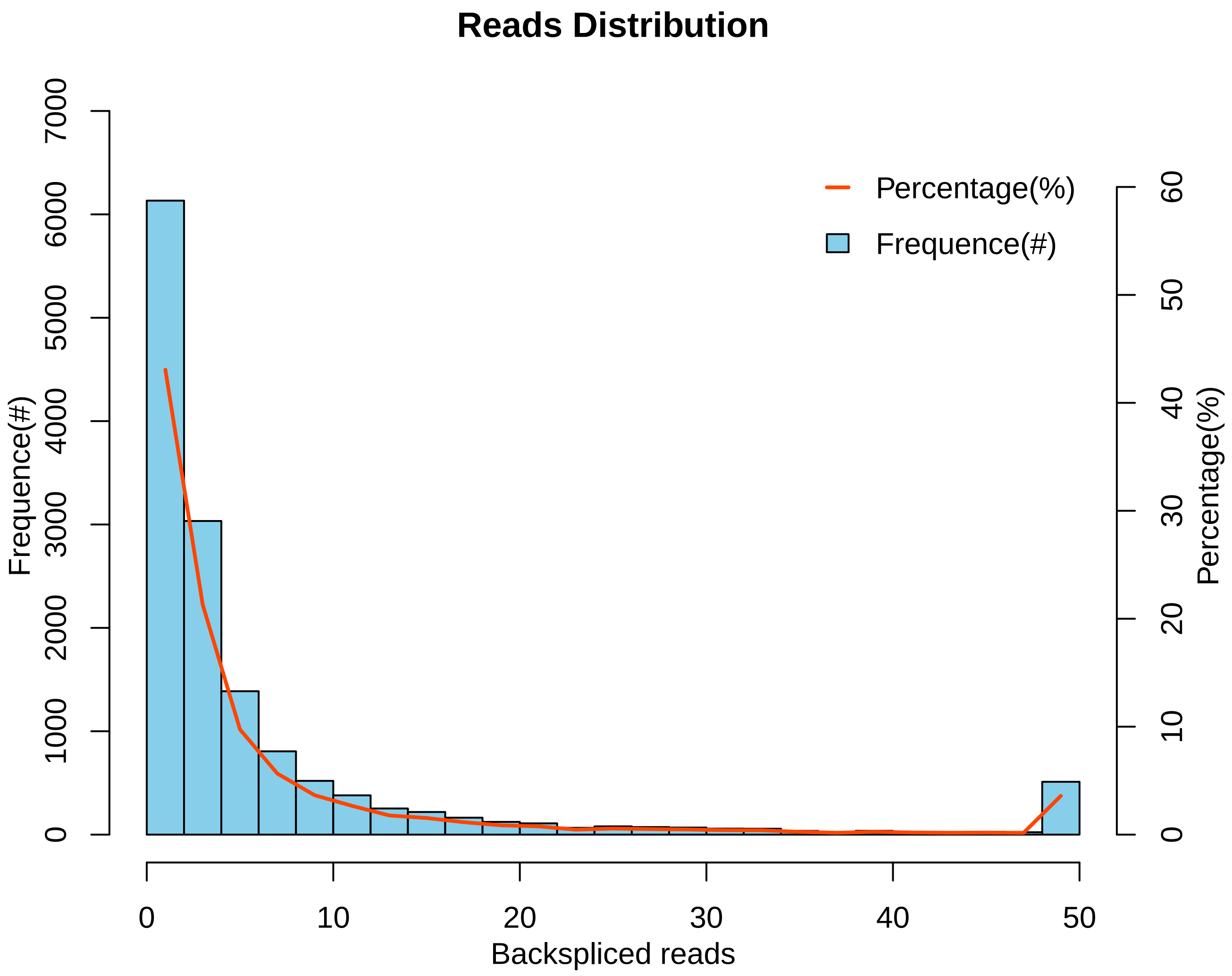

### Additional file 4

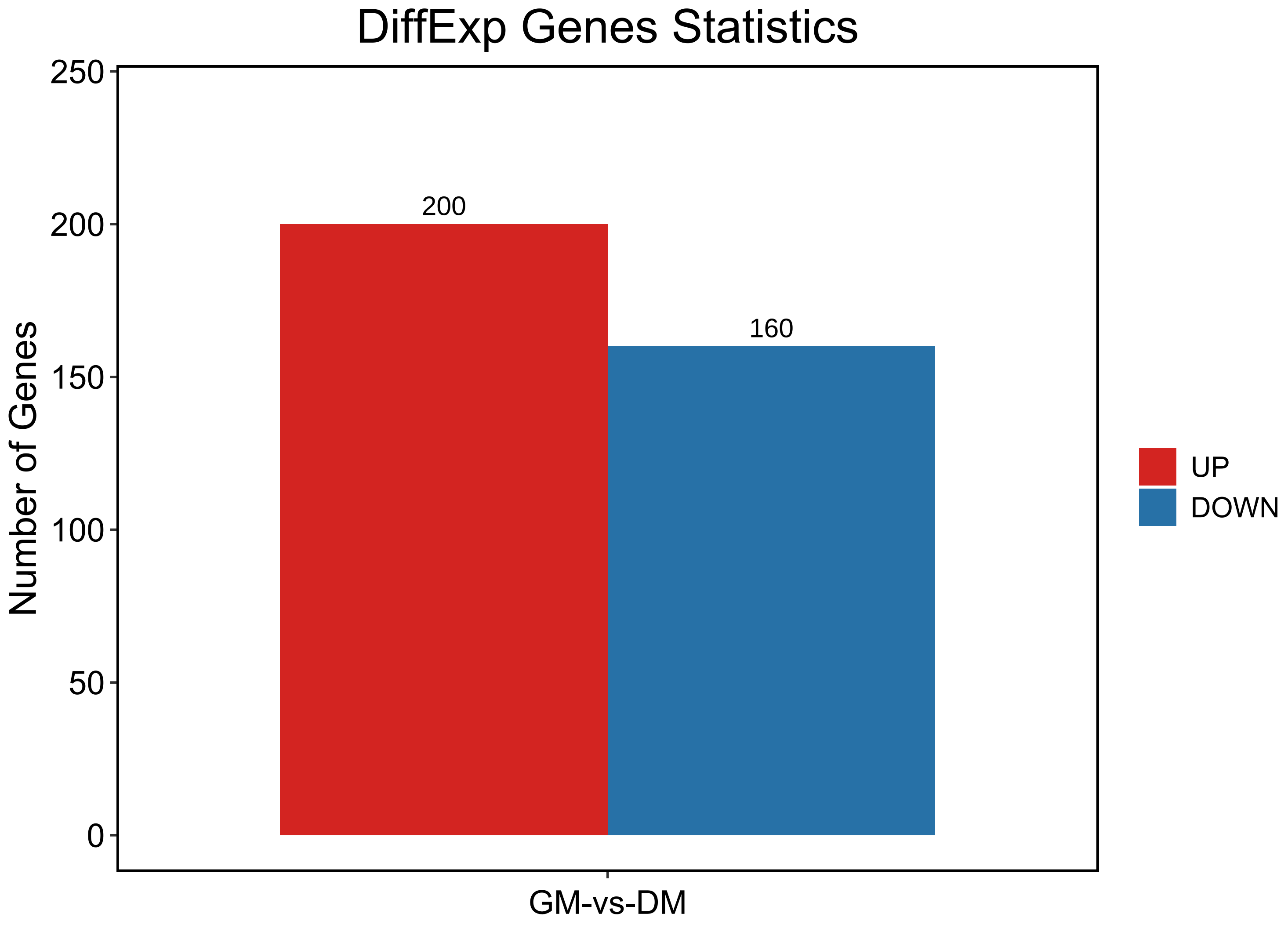

### Additional file 5

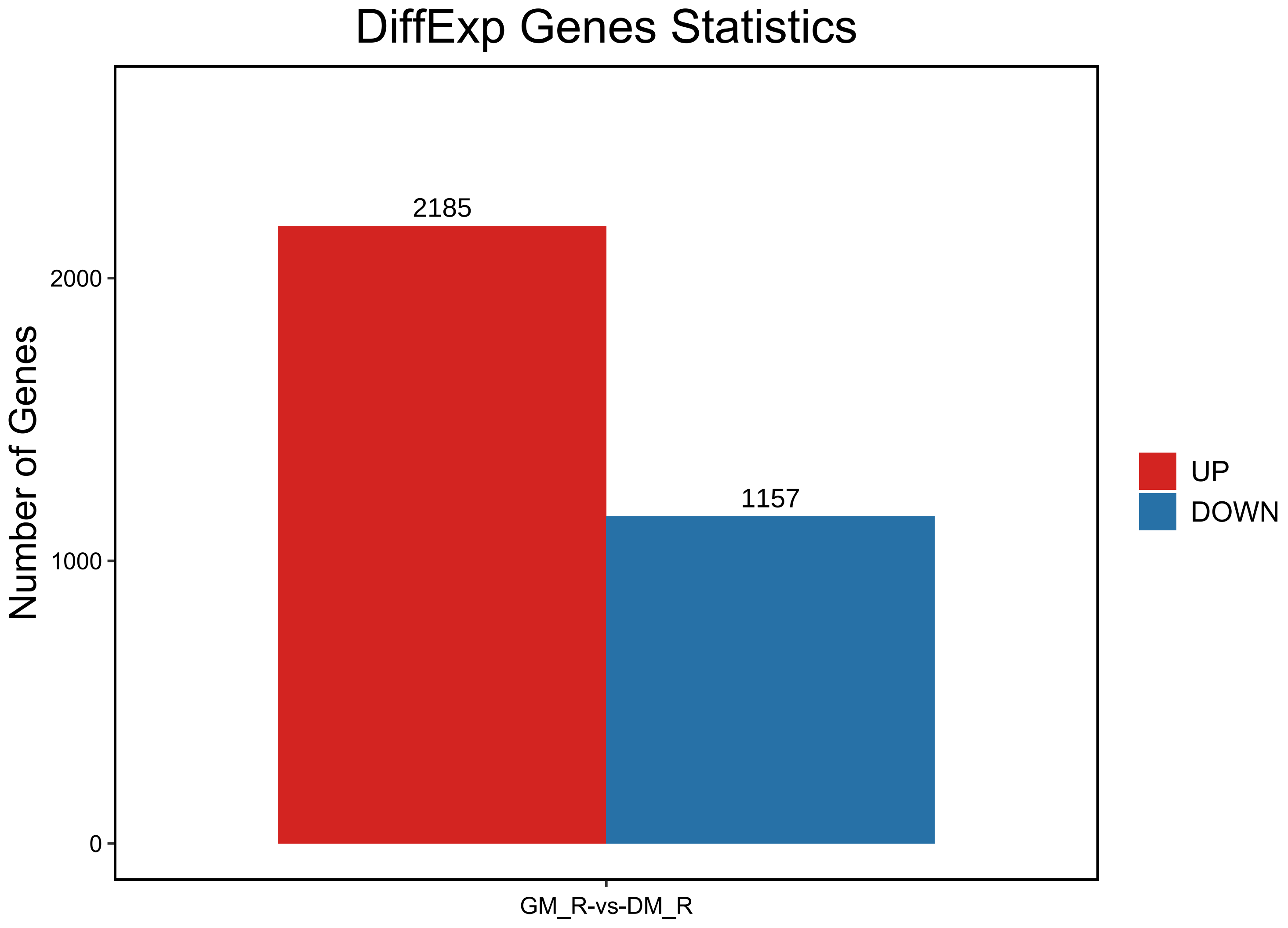
